# Contribution of rat insular cortex to stimulus-guided action

**DOI:** 10.1101/2024.10.10.617625

**Authors:** Yacine Tensaouti, Louis Morel, Shauna L. Parkes

## Abstract

Anticipating rewards is fundamental for decision-making. Animals often use cues to assess reward availability and to make predictions about future outcomes. The gustatory region of the insular cortex (IC), the so-called gustatory cortex, has a well-established role in the representation of predictive cues, such that IC neurons encode both a general form of outcome expectation as well as anticipatory outcome-specific knowledge. Here, we used Pavlovian-instrumental transfer (PIT) in male rats to assess if the IC is also required for predictive cues to exert both a general and specific influence over instrumental actions. Chemogenetic inhibition of IC abolished the ability of a reward-predictive stimulus to energize instrumental responding for reward. This deficit in general transfer was evident whether the same or different outcomes were used in the Pavlovian and instrumental conditioning phases. We observed a similar deficit in specific PIT, such that rats with IC inhibition failed to use a reward-predictive stimulus to guide choice toward actions that deliver the same food reward. Finally, we show that rats with IC inhibition also fail to show outcome-selective reinstatement. Together, these data suggest a crucial role for IC in the representation of appetitive outcomes, and particularly in using this representation to guide instrumental action.

**SIGNIFICANCE STATEMENT:** Animals frequently use cues to infer the availability of rewards and to make predictions about future outcomes. The influence of these predictive cues on behaviour can be studied using Pavlovian-instrumental transfer (PIT), in which Pavlovian outcome expectancies can elevate (general PIT) or selectively guide (specific PIT) instrumental actions. In the current study, we show that chemogenetic inhibition of the gustatory region of insular cortex (IC) abolishes both general and specific transfer, as well as the selectivity of outcome-induced reinstatement. These results demonstrate a critical role for the IC in the representation of appetitive outcomes and significantly contribute to a broader understanding of the cortical bases of PIT.

## INTRODUCTION

Environmental cues are often used to infer the availability of rewards and to make predictions about future outcomes. Our behaviour can thus be shaped by the presence of stimuli that we have learned signal particular outcomes (Colwill and Rescorla, 1988). This influence of predictive stimuli on decision-making is exemplified by Pavlovian-instrumental transfer or PIT (Walker, 1942; Estes, 1943; Colwill and Rescorla, 1988; Holmes et al., 2010), which refers to the capacity of Pavlovian outcome expectancies to affect instrumental action. Two forms of transfer have been observed. General transfer reveals that a reward-predictive stimulus elevates responding on an action that earns the same (or similar) rewarding outcome (Estes, 1943, 1948; Rescorla and Solomon, 1967), due to an increase in motivation via shared appetitive outcome features (Corbit and Balleine, 2005). In specific transfer, the predictive stimulus selectively facilitates performance on an action that delivers the same reward as the stimulus, but not other actions earning different rewards, on the basis of a shared, detailed outcome representation (Kruse et al., 1983). Thus, Pavlovian stimuli can elevate (general PIT) or selectively guide (specific PIT) instrumental action.

The gustatory region of insular cortex (IC), the gustatory cortex, has a well-documented role in the representation of predictive stimuli. In an elegant series of studies, Fontanini and colleagues (e.g., Maffei et al., 2012; Samuelsen et al., 2012, 2013; Gardner and Fontanini, 2014; Kusumoto-Yoshida et al., 2015; Vincis and Fontanini, 2016; Vincis et al., 2020) demonstrated that IC neurons show robust responses not only to auditory stimuli predicting the general availability of taste (Samuelsen et al., 2012, 2013) but also to stimuli signaling specific taste outcomes (Gardner and Fontanini, 2014; Vincis et al., 2020; see also Saddoris et al., 2009). These anticipatory IC signals are acquired with learning and extinguish, suggesting that they track the predictive status of Pavlovian stimuli (Gardner and Fontanini, 2014).

While the neural substrates of PIT have been extensively studied (see Cartoni et al., 2016), the potential involvement of the IC in transfer remains unknown. This is surprising given the clear evidence that the gustatory region of IC encodes information about anticipated outcomes, including outcome value, to guide instrumental action (Balleine and Dickinson, 2000; Saddoris et al., 2009; Parkes and Balleine, 2013; Parkes et al., 2015, 2018). The IC is also anatomically well-connected to several regions known to mediate both general and specific PIT, including the nucleus accumbens (Allen et al., 1991), mediodorsal thalamus (Wright and Groenewegen, 1996), ventral tegmental area (Ohara et al., 2003), orbitofrontal cortex (Barreiros et al., 2021; Chen et al., 2022; Mathiasen et al., 2023), and the basolateral and central nucleus of the amygdala (Sripanidkulchai et al., 1984; Yamamoto et al., 1997; Yamamoto, 2006; Gehrlach et al., 2020). Thus, the IC appears ideally placed to contribute to PIT.

Here, we examined the functional involvement of the gustatory region of the IC in general and specific PIT. We first show that general transfer is abolished under chemogenetic inhibition of IC when the same outcome is used during Pavlovian and instrumental conditioning. However, while it is largely accepted that such experimental designs indeed elicit general transfer (Hall et al., 2001; Holland and Gallagher, 2003; Corbit and Balleine, 2005, 2011), it has been noted that these “single outcome” designs might allow for mixed transfer effects (Cartoni et al., 2016). Therefore, to ensure that our results reflect deficits in general and not specific PIT, we replicated these findings using two distinct outcomes for Pavlovian and instrumental conditioning and again found that IC inhibition impaired general PIT. We then demonstrate that the IC is also required for specific PIT, as well as outcome-selective reinstatement. Together, these results indicate that IC plays a critical role in the representation of appetitive outcomes and significantly contribute to our understanding of the cortical bases of PIT.

## MATERIALS AND METHODS

### Subjects

Subjects were male Long-Evans rats, aged 3-4 months (Janvier, France). Rats were housed in pairs within plastic boxes in a climate-controlled room, maintained on a 12-hour light/dark cycle. All behavioural observations occurred during the light phase of this cycle. Rats were handled daily for five days prior to the behavioural procedures.

Two days before behaviour, rats were food restricted to maintain their weight at approximately 90-95% of their ad libitum feeding weight. All experimental protocols adhered to French (Council Directive 2013-118, February 1, 2013) and European (Directive 2010-63, September 22, 2010, European Community) legislations and received approval from the local ethics committee.

### Viral vectors

An adeno-associated viral vector carrying the inhibitory hM4Di designer receptor exclusively activated by designer drugs (DREADDs; Armbruster et al., 2007; Rogan and Roth, 2011) was obtained from Addgene (AAV8-CaMKIIa-hM4D(Gi)-mCherry; 1.46×10^13^ gc/ml; Addgene plasmid #50477; http://n2t.net/addgene:50477; RRID:Addgene_50477; gifted from Bryan Roth). A control vector lacking the hM4Di receptor was also used (AAV8-CaMKII-EGFP; 2.1×10^13^ gc/ml Addgene viral prep #50469-AAV8; http://n2t.net/addgene:50469; RRID:Addgene_50469; gifted from Bryan Roth). To activate the inhibitory receptor, the exogenous ligand deschloroclozapine (DCZ; MedChemExpress HY-42110) was dissolved in dimethyl sulfoxide (DMSO) to obtain a final concentration of 50 mg/mL, which was aliquoted and stored at −80°C (stock solution). This stock solution was further diluted in physiological saline to obtain a final concentration of 0.1 mg/mL, which was injected at 1 mL/kg (i.e., 0.1 mg/kg). DCZ was always handled in dim light conditions and fresh solutions were prepared each day. Rats were injected with DCZ or vehicle (0.2% DMSO in physiological saline) intraperitoneally (i.p.) 30 minutes before behaviour. We, and others, have previously demonstrated the efficacy of this ligand (Nagai et al., 2020; Nentwig et al., 2022; Cerpa et al., 2023).

### Surgery

Rats were anesthetized with isoflurane (5% induction, 1-2% maintenance) and placed on a stereotaxic apparatus (Kopf). The incision site was disinfected with betadine and subcutaneously injected with a local anesthetic mixture of lidocaine and ropivacaine. Vaseline was applied to the eyes and a heating pad was placed under the rat. Rats were rehydrated with subcutaneous injections of warm saline (0.9%, 10 ml/kg/hour) throughout the surgery. The viral vector was injected using a 10 µl Hamilton syringe connected to a microinjector (UMP3 UltraMicroPump II with Micro4 Controller, World Precision Instruments). 1 µl of AAV was injected at a rate of 0.2 µl/min at two sites in each hemisphere of IC. The coordinates were (in mm from bregma): AP +0.3, ML ±5.5, DV −7.3, and AP +1.3, ML ±5.5, DV −7.3 (Paxinos and Watson, 2014). After surgery, rats were subcutaneously injected with a nonsteroidal anti-inflammatory drug (meloxicam, 2 mg/kg), and sutures were covered with aluminum spray. Rats were then individually housed in a warm cage with facilitated access to food and water, and continuously monitored for 2 hours. Finally, rats were returned to their collective home-cage for post-operative care and were allowed a minimum of 8 days to recover before behavioural procedures began. Injections of vehicle or DCZ occurred 4-5 weeks after surgery.

### Behavioural Apparatus

Behavioural procedures were conducted in 8 operant cages (40 cm width × 30 cm depth × 35 cm height, Imetronic, France) that were individually enclosed in sound-resistant shells. Each chamber was equipped with two pellet dispensers that delivered 45 mg grain (BioServ) or sugar pellets (Test Diet) into a food port. Cages also contained two retractable levers that could be inserted to the left and right of the food port, speakers that provided auditory stimuli (3kHz tone or 10Hz clicker), and a house light.

### Behavioural Procedures

#### Experiment 1a: General Pavlovian-instrumental transfer – single outcome

Rats first received a habituation session to the two conditioned stimuli (CS+ and CS-, tone and clicker, counterbalanced). This session was identical to the Pavlovian conditioning sessions except no outcomes were delivered. Pavlovian conditioning began the following day. Rats received 8 Pavlovian training sessions, each lasting 1.5 h. During each session, rats were presented with 2 min presentations of the two CSs, one paired with grain pellets (CS+) and the other with no outcome (CS-). Grain pellets were delivered on a random time 30 s schedule throughout the CS+ (i.e., 4 pellets per CS+ presentation). During each session, the CS+ was presented 9 times and the CS-was presented 3 times in a pseudorandom order with a variable inter-trial interval (ITI) that averaged to 5 min (ranging from 3-7 min). Food port entries were recorded during CS+, CS-, and a 2 min interval before each CS presentation (preCS).

Following Pavlovian conditioning, rats underwent 12 sessions of instrumental training (2 sessions per day). During each session, the left and right levers were available and responding on one lever (active lever) earned grain pellets (i.e., the same outcome used for Pavlovian conditioning), whereas responding on the other lever (inactive lever) was unrewarded. Responding on the active lever was continuously reinforced on the first session and then shifted to a variable interval 30 s schedule of reinforcement for sessions 2-12. The identity of the active and inactive lever (left versus right) was counterbalanced. Rats could earn a maximum of 30 pellets per session and the maximum session duration was 30 min.

The next day, all rats underwent instrumental extinction during which the two levers were extended for 30 min and no outcomes were delivered. Rats were then given a Pavlovian extinction session, which was identical to the CS habituation session previously described. The aim of these extinction sessions was to (1) reduce the baseline rate of instrumental responding, and (2) reduce competition between the Pavlovian and instrumental responses (Holmes et al., 2010).

Twenty-four hours after the extinction sessions, rats were injected i.p. with vehicle or DCZ and, 30 min later, they were given a PIT test. At the beginning of the test, the two CSs were each presented twice in the absence of the levers. Then, both levers were extended and instrumental responding was extinguished for 8 min (in the absence of the CSs) to reduce baseline instrumental responding. Subsequently, each CS was presented twice in a pseudorandom order while both levers were extended. Each CS lasted 2 min with a fixed 4 min ITI. Food port entries and lever pressing were recorded throughout the session and responses were separated into preCS, CS+, and CS-periods (2 min each). No outcomes were delivered during the PIT test.

#### Experiment 1b: General Pavlovian-instrumental transfer – two outcomes

The procedure was identical to that described for Experiment 1a except that two distinct outcomes were used for Pavlovian conditioning and instrumental training. That is, half of the rats were trained with sugar pellets during the Pavlovian phase and grain pellets during the instrumental phase and vice versa for the remaining half. For all rats, CS+ was the clicker and CS-was the tone, and all rats were injected with DCZ prior to the general PIT test.

#### Experiment 2: Specific Pavlovian-instrumental transfer

Rats first received a single habituation session to the two CSs (tone and clicker). During this 90 min session, each CS was presented 6 times in a pseudo-random order, during which no outcomes were delivered. The duration of each CS was 2 min and the average ITI was 5 min (range 3-7 min). Then, all rats received 8 sessions of Pavlovian conditioning (one session per day) during which each CS (tone and clicker) was associated with one of the food outcomes (grain or sugar pellets, counterbalanced).

Each session consisted of four tone and four clicker presentations. During each 2 min CS, the associated reward was delivered on a 30 s random-time schedule, resulting in 4 pellet deliveries during each CS presentation. CSs were delivered pseudo-randomly with an average ITI of 5 min (range 3-7 min). Food port entries were recorded and separated into CS period and a 2 min period before each CS presentation (preCS).

Rats then received instrumental training to perform two actions to earn two distinct food rewards (e.g., left lever earns a grain pellet and right lever earns a sugar pellet, or vice versa). These food rewards were the same outcomes that were used for Pavlovian conditioning. Rats were trained under continuous reinforcement (CRF) for the first 3 days, then under a random-ratio 5 schedule (RR5) on days 4-6 and, finally, an RR10 schedule on days 7-9. During these sessions, each lever was presented twice for a maximum of 10 min each or until 20 outcomes were earned. The ITI between lever presentations was 2.5 min. Hence, each subject could earn a maximum of 40 grain and 40 sugar pellets during each session.

The next day, rats were given an instrumental extinction session followed by a Pavlovian extinction session, as described above for the general PIT paradigm. Twenty-four hours later, all rats were injected with DCZ and were then given an outcome-specific PIT test 30 min later. The test procedure was identical to the general PIT test.

Performance was evaluated under three conditions: lever pressing during the preCS period, pressing on the lever that shared the same outcome as the presented cue (same condition), and pressing on the lever that shared a different outcome with the presented cue (different condition). No outcomes were delivered during the PIT test.

#### Experiment 3: Outcome-induced reinstatement

Rats were first given instrumental training, as described for Experiment 2. The training lasted for 7 days with continuous reinforcement (CRF) for the first 3 days, then a random-ratio 5 schedule (RR5) on days 4-5 and, finally, an RR10 schedule on days 6-7. On day 8, rats were given an outcome-induced reinstatement test. The test began with a 15 min extinction period to reduce baseline instrumental responding. During this period, both levers where available but no outcomes were delivered. This was followed by 4 reinstatement trials with an ITI of 7 min. Each trial consisted of a single outcome delivery (either the grain or sugar pellet) in the following order: grain, sugar, sugar, grain.

Responding was measured during the 2 min periods immediately before (Pre) and after (Post) each outcome delivery (Ostlund and Balleine, 2007; Bradfield et al., 2015, 2018; Abiero et al., 2022). Responding during the reinstatement test was not rewarded. Rats were then given one day of rest and received a second reinstatement test on day 9 with the order of outcome deliveries counterbalanced: sugar, grain, grain, sugar. Data are presented as an average of these two tests.

Finally, rats were given an additional RR10 retraining session and then received a reinstatement test with a novel outcome (chocolate pellet) that had not been associated to either action. This session was identical to that described, with 4 chocolate pellets delivered with a fixed ITI of 7 min.

### Tissue Processing and Immunohistochemistry

After behavioural testing, rats received an i.p. injection of xylazine (20 mg/kg, i.p.) followed by an i.p. injection of pentobarbital (Euthasol; 200 mg/kg, diluted in saline). Rats were then perfused transcardially with 4% paraformaldehyde in 0.1M phosphate buffer. Subsequently, 40 µm coronal sections were cut using a VT1200S Vibratome (Leica Microsystems). Every fourth section was collected to form a series and immunoreactivity was performed for mCherry (hM4Di) but not for GFP as the endogenous fluorescence was sufficient to detect infected cells.

For hM4Di injected rats, free-floating sections were rinsed several times in 0.1M phosphate buffered saline (PBS), permeabilized with 0.3% Triton-X in 0.1M PBS (PBS-T), blocked for 1h (blocking solution: 4% normal goat serum in PBS-T) and placed in 1:1000 rabbit anti-RFP (CliniSciences, PM005) at room temperature for 24 h. Sections were then washed in PBS-T and incubated in 1:500 biotin goat-anti rabbit (Jackson ImmunoResearch, 111-065-144) diluted in PBS-T for 2 h at room temperature. Sections were then rinsed with PBS and incubated with 1:400 A594-Streptavidin (Jackson ImmunoResearch, 016-580-084) in PBS. Finally, sections were rinsed in PB 0.1M then mounted and cover-slipped with Fluoroshield with DAPI (Invitrogen, 00-4959-52).

Slides were scanned using an upright fluorescent microscope system (Leica DM5500B) attached to a Jenoptik ProgRes MF Cool (#D-07739 Jena) that was equipped with a motorized stage (Märzhäuser #W215885) driven by the open source acquisition software Micro Manager (MMStudio V1.4.10). Acquired mosaics (10x objective) were stitched to reconstruct images, using the “Stitching - Grid/Collection” plugin of the Fiji freeware (Preibisch et al., 2009). Manual tracing of injection sites for each subject was performed using the mosaic images. These hand-traced injection sites were then superimposed onto corresponding atlas sections (Paxinos & Watson, 2014) and combined to visually convey the degree of overlap in virus expression across rats.

### Experimental Design and Statistical Analyses

All experiments used a mixed method design. In Experiment 1a, separate statistical analyses were conducted on rats injected with the GFP control virus and rats injected with the inhibitory hM4Di virus. As such, the between-subject factor was treatment (vehicle versus DCZ) and within-subject factors were lever (active versus inactive), CS presentation (CS+ versus CS-), and session/time. The final group sizes, after histology and outlier exclusions, were: group GFP vehicle: n = 14, group GFP DCZ: n = 15, group hM4Di vehicle: n = 11, group hM4Di DCZ n = 11.

In Experiment 1b, the between-subject factor was group (GFP: n = 7 versus hM4Di: n = 8) and within-subject factors were lever (active versus inactive), CS presentation (CS+ versus CS-), and session/time. All rats were injected with DCZ. For Experiment 2, the between-subject factor was again group (GFP: n = 13 versus hM4Di: n = 12) and within-subject factors were CS presentation (same versus different) and session/time, and all rats were injected with DCZ. Finally, in Experiment 3, the between-subject factor was group (GFP: n = 12 versus hM4Di: n = 12) and the within-subject factors were lever (reinstated or non-reinstated), period (pre-versus post-outcome delivery) and session/time. Again, all rats were injected with DCZ.

The dependent variables were the rate of lever presses or food port entries (FPE), which are presented as the response rate during the CS presentation minus the response rate during the baseline preCS period (2 min period preceding each CS presentation) for the PIT experiments. Data were analysed using a mixed-model ANOVA followed by simple effects analyses to establish the source of any significant interactions. Statistical significance was set at p ≤ 0.05. Data are presented as mean ± SEM and individual data points are supplied on histograms. Rats were excluded if they failed to properly learn the Pavlovian associations, defined as greater responding during CS-than during CS+ during the final two days of training for general PIT or greater responding during the preCS period versus the CS period during the final two training days for specific PIT. Four rats were excluded based on this criterion, two from general PIT and two from specific PIT.

## RESULTS

### Histology

Schematics of viral expression in IC for each rat are illustrated in **Figure 1A** along with representative images (**Figure 1B**). In total, two rats were excluded due to unilateral viral expression, one in Experiment 1a and one in Experiment 2.

**Figure 1.**
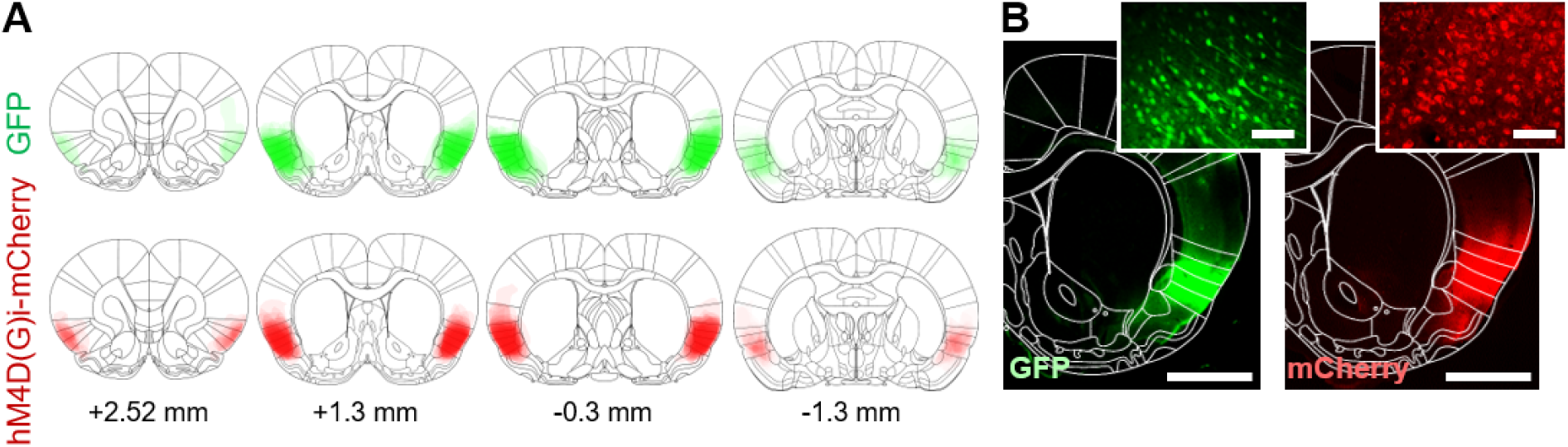
**(A)** Schematics of the viral expression in IC (in mm from bregma) for rats injected with the inhibitory DREADD (hM4Di) or control virus (GFP) for all experiments. **(B)** Representative images from a rat in group GFP and a rat in group hM4Di (mCherry). Scale bars represent 1 mm in the map image (10x) and 50 µm in the magnified inset image (20x).

### Experiment 1a: General transfer is impaired following IC inhibition

We first assessed the impact of chemogenetic inhibition of insular cortex (IC) on general PIT using a single outcome in Pavlovian conditioning and instrumental training. Rats received bilateral IC injections of a virus carrying the inhibitory DREADD (hM4Di) or a control virus (GFP). One rat was excluded due to unilateral viral infection and two rats were excluded because they failed to properly discriminate between CS+ and CS-during Pavlovian conditioning. This yielded the following between-subject group sizes: GFP vehicle: n = 14, GFP DCZ: n = 15; hM4Di vehicle: n = 11, hM4Di DCZ n = 11.

Following recovery, rats underwent the general PIT procedure with a single outcome, as shown in Figure 2A. They first received Pavlovian conditioning during which CS+ was paired with the food reward and CS-was not. Rats were then given instrumental training whereby responding on the active lever earned the food reward while responding on the inactive lever was unrewarded. The food reward was the same for Pavlovian and instrumental phases. Finally, all rats received a transfer test that assessed lever pressing in the presence of the two stimuli (CS+ and CS-). Rats were injected i.p. with vehicle or DCZ 30 min before the general PIT test.

**Figure 2.**
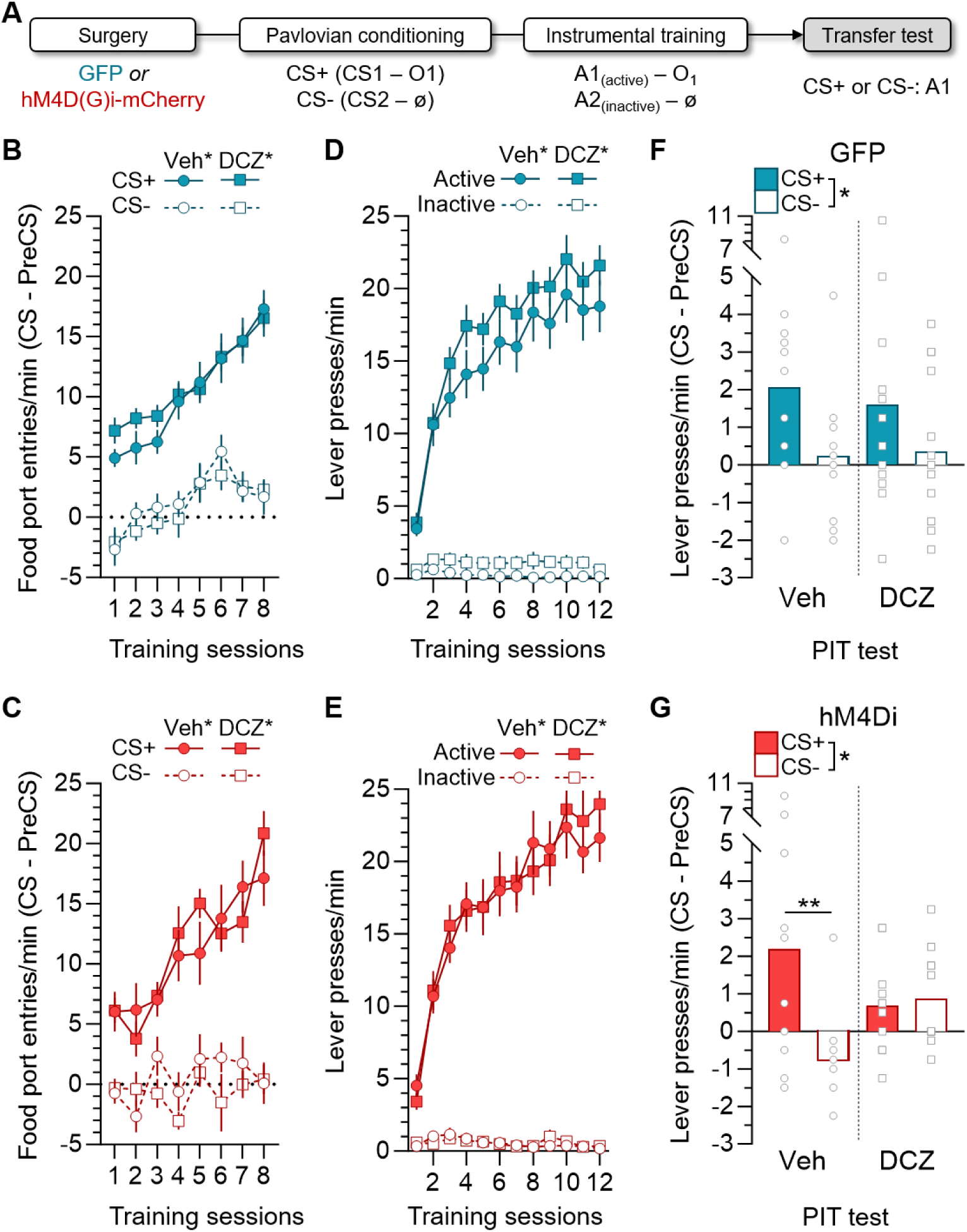
IC inhibition impairs general PIT when a single outcome is used. **(A)** Outline of behavioural procedure, vehicle or DCZ was injected during the transfer test. **(B, C)** Mean (±SEM) rate of food port entries (FPE) during CS+ (filled plot) and CS-(open plot) across Pavlovian conditioning for GFP (B) and hM4Di (C) rats. Calculated as FPE rate during the 2 min CS minus the FPE rate during the 2 min before CS onset (preCS). Vehicle (veh) groups are shown in circles and DCZ groups are in squares (veh*; DCZ* indicates treatment to be during the general PIT test). **(D, E)** Lever pressing rate (mean ±SEM) across instrumental training on active (filled plot) and inactive levers (open plot) for GFP (D) and hM4Di (E) rats. **(F, G)** Performance on active lever during CS+ (filled bars) and CS-(open bars) in the general PIT test for GFP (F) and hM4Di (G) rats treated with vehicle or DCZ. Calculated as the pressing rate during the 2 min CS minus the pressing rate during the preCS. *statistical significance.

Pavlovian conditioning proceeded smoothly for all groups. For GFP-injected rats (Figure 2B), there was a main effect of CS (F_1,27_ = 241.8, p < 0.01), indicating that the rate of food port entries was higher in the presence of the rewarded stimulus (CS+) than in the presence of the unrewarded stimulus (CS-). There was also a main effect of session (F_1,27_ = 180.07, p < 0.01), but no effect of treatment to be or any significant interactions (largest F_1,27_ = 1.32, p = 0.26). A similar pattern of results was observed for rats injected with the hM4Di virus (Figure 2C), with a main effect of CS (F_1,20_ = 206.48, p < 0.01) and session (F_1,20_ = 33.15, p < 0.01), but no effect of treatment to be or significant interactions (largest F_1,20_ = 1.03, p = 0.32).

Rats also successfully acquired instrumental responding with a main effect of session (F_1,27_ = 157.87, p < 0.01), lever (F_1,27_ = 264.91, p < 0.01), and a lever x session interaction (F_1,27_ = 200.92, p < 0.01) detected for GFP (Figure 2D) and hM4Di groups (Figure 2E; session: F_1,20_ = 160.51, p < 0.01; lever: F_1,20_ = 291.50, p < 0.01; session × lever interaction: F_1,20_ = 148.94, p < 0.01). Simple effect analyses conducted on the session x lever interactions indicated that lever pressing increased on the active (GFP: F_1,27_ = 182.22, p < 0.01; hM4Di: F_1,20_ = 159.48, p < 0.01) but not inactive levers (GFP: F_1,27_ = 2.03, p = 0.17; hM4Di: F_1,20_ = 3.75, p = 0.07). There were no significant differences in responding between rats that would receive vehicle or DCZ during the upcoming transfer test or any interactions between treatment and the other factors (GFP: largest F_1,27_ = 2.56, p = 0.12; hM4Di: largest F_1,20_ = 0.87, p = 0.36).

To reduce baseline instrumental responding and Pavlovian/instrumental response competition, all rats underwent a 30 min instrumental extinction session followed by a Pavlovian extinction session on the day preceding the transfer test (data not shown). For GFP rats, there was a significant main effect of lever (F_1,27_ = 187.95, p < 0.01), time (F_1,27_ = 141.55, p < 0.01), and a significant lever x time interaction (F_1,27_ = 94.00, p < 0.01) but no effect of treatment to be (F_1,27_ = 0.34, p = 0.57). Simple effects analyses indicated that responding on both the active (F_1,27_ = 118.11, p < 0.01) and inactive lever (F_1,27_ = 18.58, p < 0.01) decreased across the session, although this decrease was greater for the active lever. Similar results were found for hM4Di rats (main effect of lever: F_1,20_ = 243.24, p < 0.01; time: F_1,20_ = 93.74, p < 0.01; lever x time interaction: F_1,20_ = 76.58, p < 0.01; no effect of treatment to be: F_1,20_ = 0.58, p = 0.46), and responding on both the active (F_1,20_ = 98.20, p < 0.01) and inactive lever (F_1,20_ = 5.24, p < 0.01) decreased across the session. The rate of food port entries across the Pavlovian extinction session did not differ between CS+ and CS-for GFP-injected rats (F_1,27_ = 0.75, p = 0.39), however hM4Di-injected rats responded more during CS+ than CS-(F_1,20_ = 7.43, p = 0.01). There were no main effects of time or treatment to be for GFP (largest F_1,27_ = 0.66, p = 0.42) or hM4Di rats (largest F_1,20_ = 2.32, p = 0.14), and no interactions (GFP: largest F_1,27_ = 2.94, p = 0.10; hM4Di: largest F_1,20_ = 0.8, p = 0.38).

Performance on the active lever during the general PIT test is shown in Figure 2F for GFP- and Figure 2G for hM4Di-injected rats. For GFP rats, lever pressing was higher during the CS+ than during the CS-(F_1,27_ = 5.25, p = 0.03), and there was no effect of treatment (F_1,27_ = 0.09, p = 0.77) or stimulus × treatment interaction (F_1,27_ = 0.2, p = 0.66), indicating successful general transfer.

A different pattern of results was observed for hM4Di rats. Statistical analyses revealed no main effect of treatment (F_1,20_ = 0.01, p = 0.91), but a main effect of CS (F_1,20_ = 4.6, p = 0.04) and a significant CS x treatment interaction (F_1,20_ = 5.86, p = 0.03). Simple effect analyses indicated that lever pressing was higher during CS+ than during CS-for rats that received vehicle (F_1,20_ = 7.31, p = 0.01) but not for rats that received DCZ (F_1,20_ = 0.7, p = 0.41).

We also examined the rate of food port entries during the transfer test (Figure 3A) and found that both GFP (F_1,27_ = 8.30, p < 0.01) and hM4Di rats (F_1,20_ = 10.64, p < 0.36) entered the food port more during CS+ than CS-, and there was no effect of treatment (GFP: F_1,27_ = 0.47, p = 0.50; hM4Di: F_1,20_ = 0.97, p = 0.34) or CS x treatment interaction (GFP: F_1,27_ = 0.64, p = 0.43; hM4Di: F_1,20_ = 1.10, p = 0.33).

**Figure 3.**
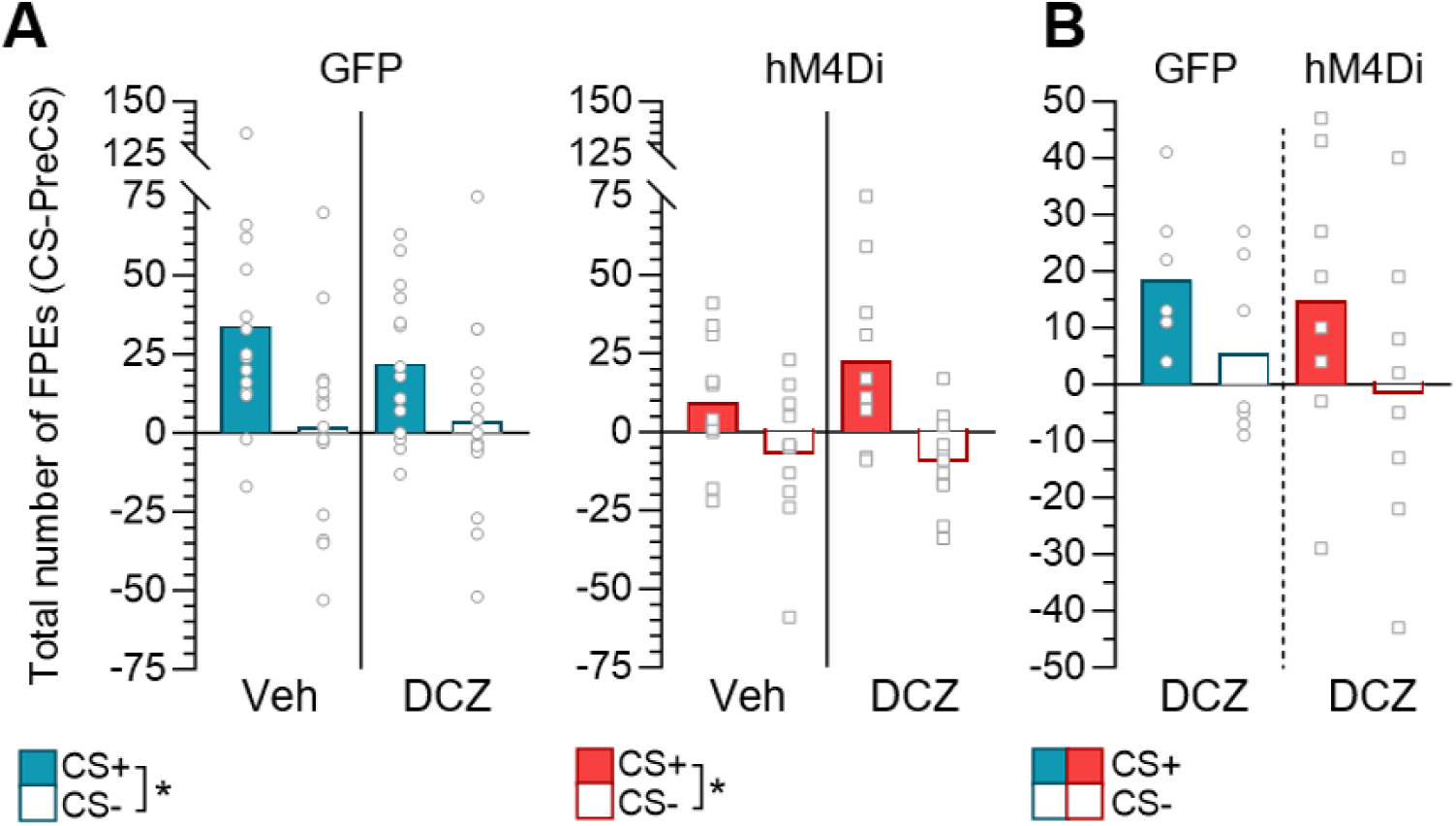
Food port entries during the general transfer tests. Mean number of food port entries (FPE) during CS+ and CS-during the transfer tests in Experiment 1a **(A)** and Experiment 1b **(B)**. Due to potential response competition between FPE and lever presses during the transfer test, we plotted and analysed total FPE during the four CS alone presentations prior to lever extension and the four CS presentations while the levers were present (i.e., 8 minutes total for CS+ and CS-). Data are presented as the total number of FPE during each CS period minus the number FPE during the 2 min period before each CS onset (preCS). *indicates statistical significance.

**Figure 4.**
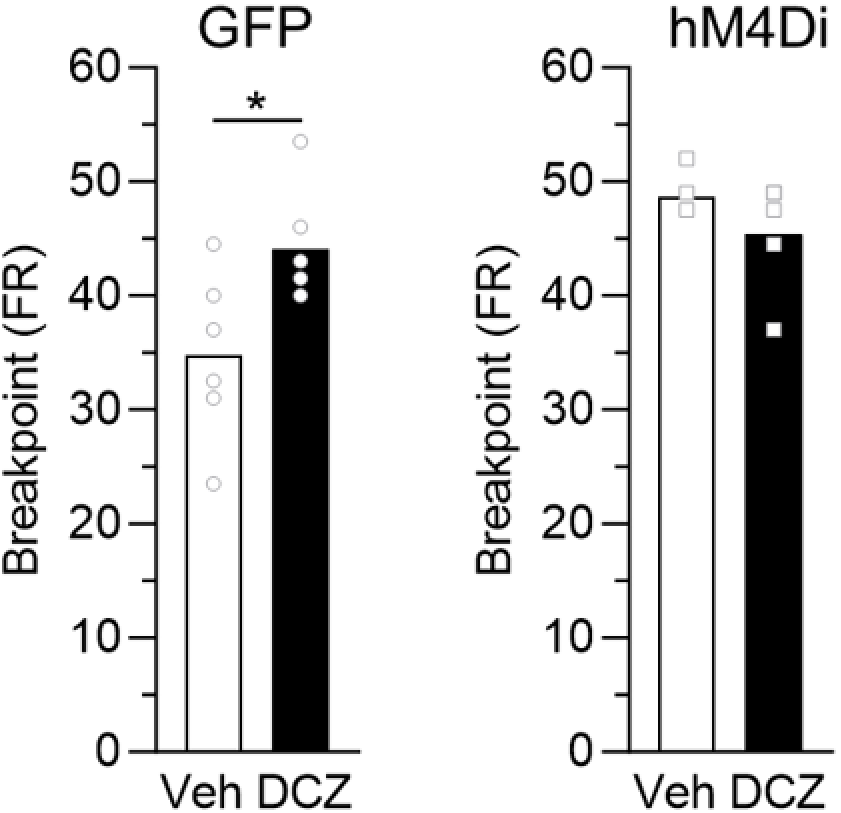
Performance in a progressive ratio task. Breakpoints, defined as the last fixed ratio (FR) completed (i.e., number of lever presses to obtain the final pellet), for GFP (left) and hM4Di rats (right) that were injected with vehicle (white bars) or DCZ (black bars) 30 min prior to the progressive ratio task.

Finally, a subset of the same rats used for general PIT underwent a progressive ratio (PR) task to assess motivation to lever press under IC inhibition (GFP vehicle n = 6; GFP DCZ n = 6; hM4Di vehicle n = 5; hM4Di DCZ n = 5). Rats were injected with vehicle or DCZ 30 minutes prior to PR. During the PR, rats were required to press the lever an increasing number of times on a fixed ratio (FR) 3 to obtain successive food pellet rewards (+3 increments, e.g., 1 press for the first pellet, 4 presses for the second pellet, 7 presses for the third pellet, etc.). The breakpoint, defined as the last fixed ratio completed (number of lever presses to obtain the final pellet), was recorded. Statistical analyses revealed that the breakpoint was significantly lower for GFP rats injected with vehicle compared to GFP rats injected with DCZ (F_1,10_ = 6.31, p = 0.03) however, the breakpoint did not differ for hM4Di rats injected with vehicle versus those injected with DCZ (F_1,8_ = 1.86, p = 0.21).

### Experiment 1b: IC inhibition also impairs general transfer when two distinct outcomes are used

The procedure was identical to that described for Experiment 1a except that half of the rats were trained with sugar pellets during the Pavlovian phase and grain pellets during the instrumental phase and vice versa for the remaining half of the rats (Figure 5A). One rat was excluded for responding > 2 standard deviations from the mean during CS+ at the end of the Pavlovian extinction session (>20 entries per min during final two extinction trials). As in Experiment 1a, rats received bilateral IC injections of the inhibitory hM4Di virus (n = 8) or the control GFP virus (n = 7) and all rats were injected with DCZ 30 min prior to the general PIT test.

**Figure 5.**
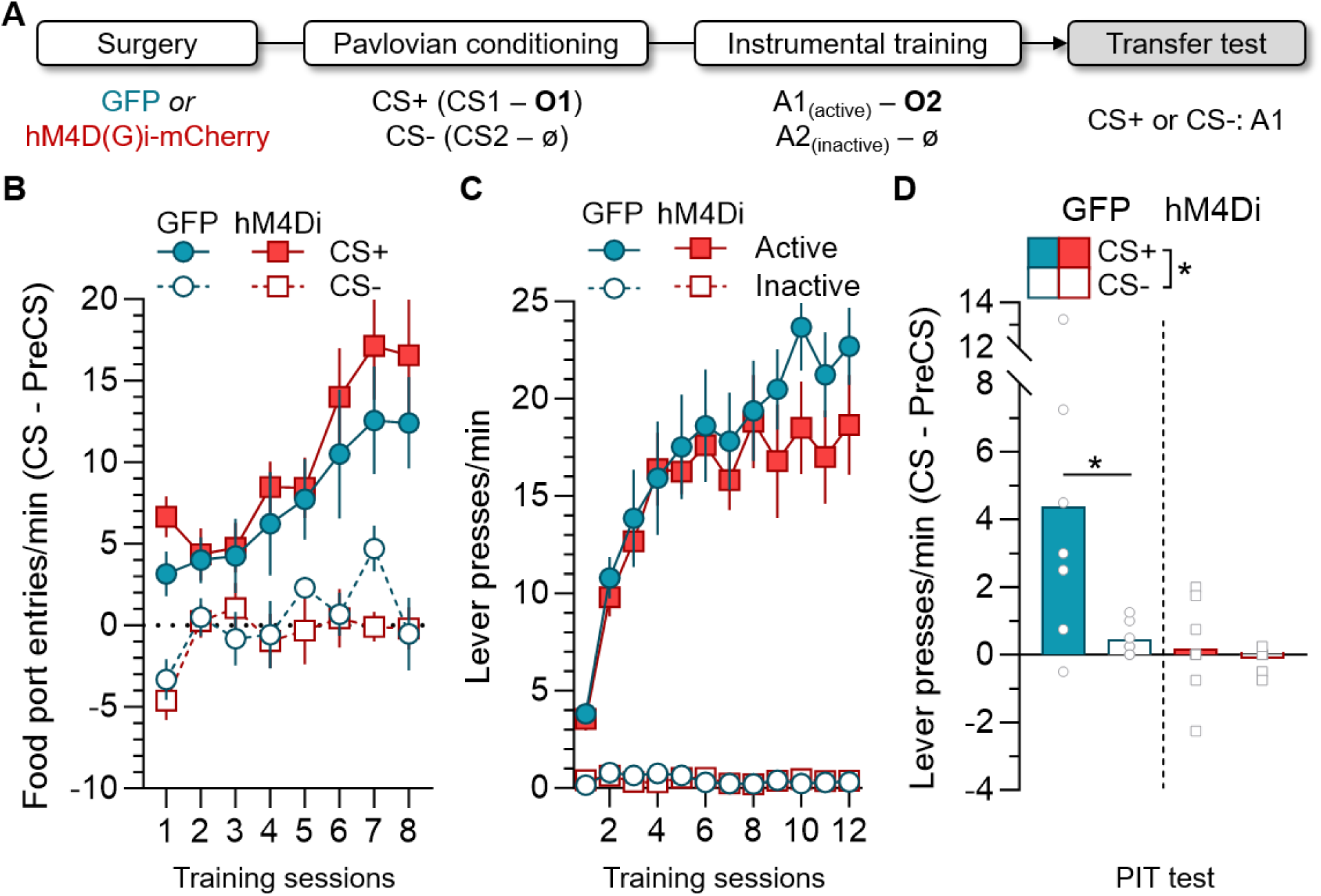
IC inhibition impairs general PIT when different Pavlovian and instrumental outcomes are used. **A.** Outline of the behavioural procedure, vehicle or DCZ was injected during the transfer test. **B.** Mean (±SEM) rate of food port entries (FPE) during CS+ (filled plots) and CS-(open plots) across Pavlovian conditioning for GFP and hM4Di rats. Calculated as the rate of entries during the 2 min CS minus the rate of entries during the 2 min before CS onset (preCS). **C.** Mean (±SEM) lever pressing rate across instrumental training on the active (filled plots) and inactive lever (open plots) for GFP and hM4Di rats. **D.** Performance on the active lever during CS+ (filled bars) and CS-(open bars) in the general PIT test for GFP and hM4Di rats under DCZ injection. Performance is calculated as the rate of pressing during the 2 min CS minus the rate of pressing during the 2 min before CS onset (preCS). *statistical significance.

Across Pavlovian conditioning (Figure 5B), food port entries were greater during CS+ than during CS-(F_1,13_ = 39.38, p < 0.01) and entries increased across training sessions (F_1,13_ = 38.27, p < 0.01) but this increase was greater for CS+ (CS x session interaction: F_1,13_ = 9.83, p < 0.01; simple effect of session for CS+: F_1,13_ = 23.15, p < 0.01; and CS-: F_1,13_ = 7.87, p = 0.02). GFP and hM4Di groups did not differ (F_1,13_ = 0.29, p = 0.60) and there were no interactions between group and any other factors (largest F_1,13_ = 1.4, p = 0.26).

Instrumental training (Figure 5C) was also successful with statistical analyses revealing a main effect of session (F_1,13_ = 56.28, p < 0.01), lever (F_1,13_ = 165.10, p < 0.01), and a lever x session interaction (F_1,13_ = 66.88, p < 0.01). Simple effects analyses revealed that responding increased on the active (F_1,13_ = 62.33, p < 0.01) but not inactive lever (F_1,13_ = 0.98, p = 0.34). There was no main effect of group or any interactions between group and the other factors (largest F_1,13_ = 2.18, p = 0.16).

Lever pressing decreased across the instrumental extinction session (data not shown), with a main effect of session (F_1,13_ = 117.67, p < 0.01), lever (F_1,13_ = 146.52, p < 0.01), and a session x lever interaction (F_1,13_ = 109.03, p < 0.01). Simple effect analyses indicated that responding decreased across sessions on the active (F_1,13_ = 125.31, p < 0.01) but not the inactive lever (F_1,13_ = 2.92, p = 0.11). No main effect of group was detected (F_1,13_ = 0.63, p = 0.44) but the three-way interaction approached significance (F_1,13_ = 4.29, p = 0.06), suggesting that the decrease in responding on the active lever was perhaps greater in group GFP than group hM4Di. During Pavlovian extinction (data not shown), food port entries were greater during CS+ than during CS-(F_1,13_ = 6.21, p = 0.03), and there was no main effect of group, session, or any significant interactions (largest F_1,13_ = 2.83, p = 0.12). Figure 5D shows responding on the active lever during the general PIT test. Statistical analyses revealed a main effect of CS (F_1,13_ = 6.62, p = 0.02) and group (F_1,13_ = 7.10, p = 0.02), indicating that pressing was greater during the CS+ and that GFP rats pressed more than hM4Di rats. Importantly, there was also a significant CS x group interaction (F_1,13_ = 4.82, p = 0.05), and simple effects analyses indicated that group GFP responded more during CS+ than CS-(F_1,13_ = 10.66, p < 0.01) but this was not the case for group hM4Di (F_1,13_ = 0.08, p = 0.78). Food port entries did not differ between groups with no main effect of CS, group, or CS x group interaction detected (largest F_1,13_ = 2.48, p = 0.14) (see Figure 3B).

### Experiment 2: Specific transfer is impaired following IC inhibition

We next examined if IC is also required for specific PIT. Rats received bilateral IC injections of the inhibitory hM4Di virus (n = 14) or the control GFP virus (n = 14). One rat was excluded due to misplaced viral expression and two rats were excluded because they failed to learn Pavlovian conditioning. This yielded the following between-subject group sizes: GFP: n = 13, hM4Di: n = 12. All rats were injected i.p. with DCZ before the specific PIT test.

Following recovery from surgery, rats underwent the specific PIT protocol (Figure 6A). They first received Pavlovian conditioning during which two distinct conditioned stimuli (CS1 and CS2) were associated with two different food outcomes (O1 and O2).

**Figure 6.**
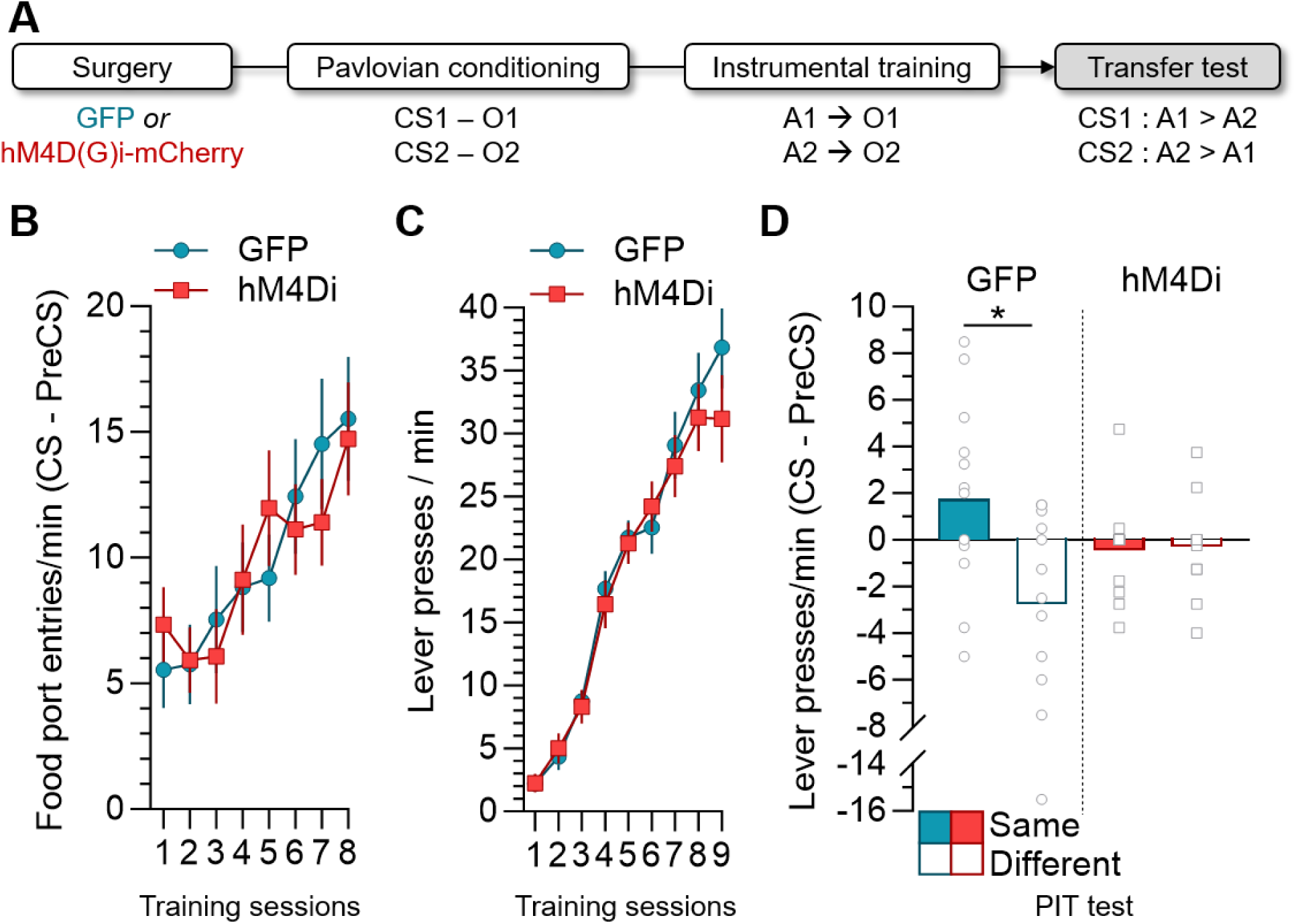
IC inhibition abolishes specific PIT. **(A)** Outline of the behavioural procedure. **(B)** Mean (±SEM) rate of food port entries (FPE) during CS presentations (clicker and tone averaged) across Pavlovian conditioning for GFP (circles) and hM4Di rats (squares). Calculated as the rate of entries during the 2 min CS minus the rate of entries during the 2 min before CS onset (preCS). **(C)** Mean (±SEM) lever pressing rate across instrumental training (left and right lever averaged) for GFP (circles) and hM4Di rats (squares). **(D)** Lever pressing during the CS that shared the same outcome (Same; filled bars) and the different outcome (Different; open bars) in the specific PIT test for GFP and hM4Di rats under DCZ injection. Performance is calculated as the rate of pressing during the 2 min CS minus the rate of pressing during the 2 min before CS onset (preCS).

**Figure 7.**
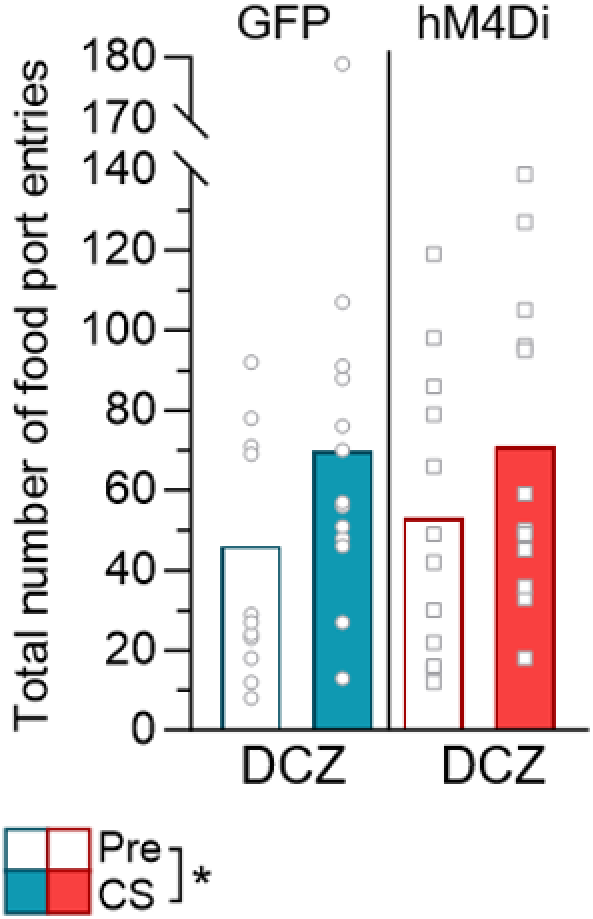
Food port entries during the specific transfer test. Mean number food port entries (FPE) during the CSs and the preCS period (Pre) during the specific transfer test. Due to potential response competition between FPE and lever presses during the transfer test, we plotted and analysed total FPE during the four CS alone presentations prior to lever extension and the four CS presentations while the levers were present. Data are presented as the total number of FPE during each CS period minus the number FPE during the 2 min period before each CS onset (preCS). *statistical significance.

Rats were then trained to perform one action (e.g., left lever) to earn one outcome (e.g., O1) and another action (e.g., right lever) to earn a different outcome (e.g., O2). Finally, all rats were injected with DCZ and then received a transfer test that assessed responding on both levers during the presence of the two stimuli (CS1 and CS2).

As shown in Figure 6B, Pavlovian conditioning was successful for both GFP and hM4Di groups with rats increasing their food port entries across sessions (F_1,23_ = 23.69, p < 0.01). There was no difference between GFP and hM4Di groups (F_1,23_ = 0.01, p = 0.92) and no significant interaction (F_1,23_ = 0.47, p = 0.50). Rats also acquired instrumental responding, with a main effect of session (F_1,23_ = 248.23, p < 0.01) but no effect of group or session x group interaction (Figure 6C; largest F_1,23_ = 0.9, p = 0.35).

All rats were then given an instrumental extinction session followed by a Pavlovian extinction session on the day preceding the transfer test (data not shown). Instrumental responding decreased across the extinction session (F_1,23_ = 133.42, p < 0.01), and there was no significant difference between groups or a significant group x session interaction (largest F_1,23_ = 2.63, p = 0.12). Food port entries across the Pavlovian extinction session did not differ between groups (F_1,23_ = 0.66, p = 0.43) and there was no effect of time or group x time interaction (largest F_1,23_ = 0.85, p = 0.37).

The results from the specific PIT test are shown in Figure 6D. Data are plotted as the rate of lever presses (collapsed across A1 and A2) during the stimulus (CS1 or CS2) that shared a common outcome (same condition) or a different outcome (different condition). Statistical analyses revealed no main effect of group (F_1,23_ = 0.03, p = 0.86) or lever (F_1,23_ = 3.96, p = 0.06) but a significant group x lever interaction (F_1,23_ = 4.5, p = 0.045). Simple effects conducted on the interaction indicated that GFP rats responded significantly more on the same lever than on the different lever (F_1,23_ = 8.81, p = 0.007) however, hM4Di rats did not (F_1,23_ = 0.008, p = 0.93).

Finally, we observed that food port entries were greater during the CSs than during the preCS periods (F_1,23_ = 11.06, p < 0.01), with no difference between groups (F_1,23_ = 0.09, p = 0.77) or CS period x group interaction (F_1,23_ = 0.21, p = 0.65). This indicates that the expression of Pavlovian conditioning was intact in both groups.

### Experiment 3: IC inhibition abolishes outcome-selective reinstatement

The previous experiments showed that IC inhibition caused deficits in both general and specific PIT. Importantly, PIT relies on the ability of a stimulus-induced outcome *representation* to energise or selectively bias instrumental action. In this final experiment, we asked if similar deficits would be observed if the outcome itself was physically present. That is, would IC inhibition also impair the ability of an appetitive outcome to reinstate responding on extinguished actions. To achieve this, we used outcome-induced reinstatement, whereby a previously extinguished instrumental action is reinstated by “free” delivery of its associated outcome (Figure 8A). Rats received bilateral IC injections of the hM4Di (n = 12) or the control GFP virus (n = 12) and no rats were excluded. All rats were injected i.p. with DCZ 30 min prior to the reinstatement test.

**Figure 8.**
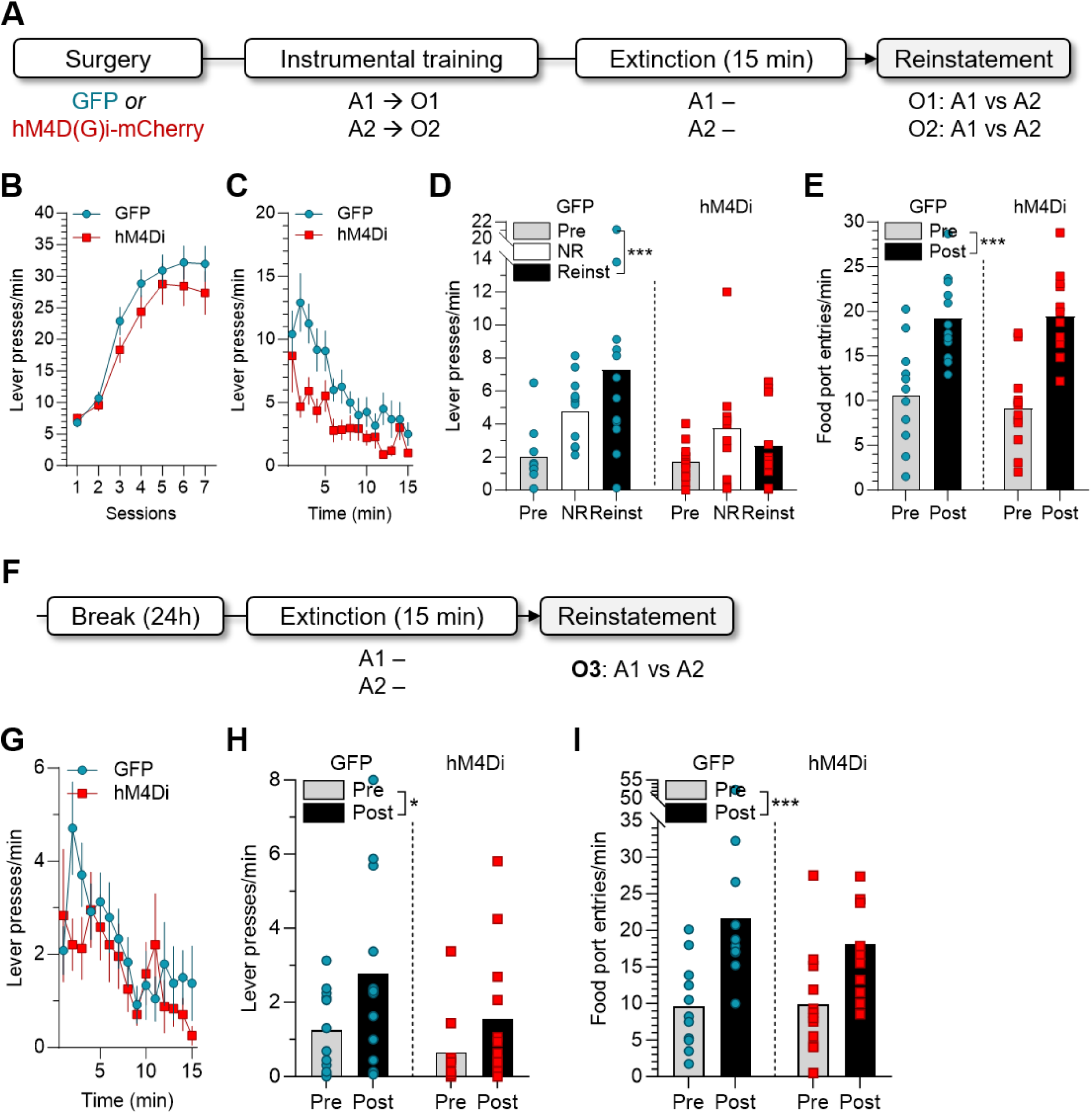
IC inhibition impairs the selectivity of outcome-induced reinstatement. **(A)** Behavioural procedure. **(B, C)** Lever pressing rate (±SEM) across instrumental training and the extinction period that preceded the reinstatement test, respectively. **(D)** Lever pressing rate during the reinstatement test before (Pre; averaged across levers) and after outcome deliveries on the non-reinstated (NR) and reinstated lever (Reinst). **(E)** Food port entry (FPE) rate pre- and post-outcome delivery during the test. **(F)** Behavioural procedure for the outcome-induced reinstatement test with a novel outcome (O3). **(G)** Lever pressing rate (±SEM) across the extinction period that preceded the reinstatement test. **(H)** Lever pressing rate pre- and post-outcome deliveries, averaged across levers. **(I)** FPE rate pre- and post-outcome deliveries.*,*** statistical significance.

Following recovery from surgery, rats received instrumental training with two A-O associations (A1-O1; A2-O2). This training proceeded smoothly (Figure 8B), with responding increasing across days (F_1,22_ = 110.70, p < 0.001) and no significant difference between groups or significant group x session interaction (largest F_1,22_ = 1.14, p = 0.30). Figure 8C shows lever pressing across the 15 min extinction session that immediately preceded the outcome-induced reinstatement test. There was a main effect of group, such that hM4Di rats pressed less than GFP rats (F_1,22_ = 15.736, p = 0.001), and a main effect of time (F_1,22_ = 62.361, p < 0.001) as well as a group x time interaction that approached significance (F_1,22_ = 4.22, p = 0.052).

The results from the outcome-induced reinstatement test are shown in Figure 8D. Overall, GFP rats responded significantly more than hM4Di rats (F_1,22_ = 4.42, p = 0.047). There was also a significant main effect of responding *before* (pre) versus *after* outcome deliveries, averaged across the non-reinstated (NR) and reinstated (Reinst) levers, such that lever pressing increased following outcome deliveries (F_1,22_ = 36.40, p < 0.001).

There was also a significant interaction with group such that the difference in instrumental responding *before* versus *after* outcome delivery was greater in the GFP group than in the hM4Di group (F_1,22_ = 7.40, p = 0.01). However, simple effect analyses confirmed that responding was indeed higher after outcome delivery for both groups (GFP: F_1,22_ = 38.31, p < 0.001; hM4Di: F_1,22_ = 5.49, p = 0.03), indicating successful (non-selective) outcome-induced reinstatement in GFP and hM4Di groups.

We then assessed if outcome-*selective* reinstatement was intact by examining the difference in responding on the non-reinstated versus reinstated lever immediately following outcome delivery. While no significant overall difference was detected (F_1,22_ = 0.77, p = 0.39), there was a significant interaction with group (F_1,22_ = 4.43, p = 0.047). Simple effect analyses indicated that group GFP responded more on the reinstated than the non-reinstated lever (F_1,22_ = 4.45, p = 0.047), but hM4Di rats did not (F_1,22_ = 0.76, p = 0.39). Thus, while both GFP and hM4Di rats showed non-selective outcome-induced reinstatement (albeit less so in hM4Di rats), only GFP rats showed outcome-selective reinstatement. Food port entries (Figure 8E) during the 2 min period after outcome deliveries were greater than during the 2 min prior to outcome deliveries (F_1,22_ = 78.95, p < 0.001), with no difference between groups or significant period x group interaction (largest F_1,22_ = 0.61, p = 0.44).

Finally, to confirm that IC inhibition did indeed leave non-selective outcome-induced reinstatement intact, rats received one session of instrumental retraining and then underwent an outcome-induced reinstatement test with a novel food reward (O3; Figure 8F). During the extinction period (Figure 8G), overall responding significantly decreased (F_1,22_ = 29.25, p < 0.001) and there was no main effect of group or group x time interaction (largest F_1,22_ = 0.87, p = 0.36). The outcome-induced reinstatement results are shown in Figure 8H, averaged across the two levers. There was a main effect of period (F_1,22_ = 7.76, p = 0.01), indicating that lever pressing was greater post outcome delivery and there was no effect of group or group x period interaction (largest F_1,22_ = 2.59, p = 0.12). Thus, the novel food reward successfully reinstated instrumental responding in both GFP and hM4Di rats. Food port entries (Figure 8I) were again greater during the 2 min period after outcome deliveries than during the 2 min prior to outcome deliveries (F_1,22_ = 23.74, p < 0.001), with no difference between groups or significant period x group interaction (largest F_1,22_ = 0.85, p = 0.37).

## DISCUSSION

Here, we examined the role of the gustatory region of the insular cortex (IC) in the general and specific forms of Pavlovian-instrumental transfer (PIT). Our findings revealed that chemogenetic inhibition of the IC during the transfer test abolished both general and specific PIT, indicating that IC is required for predictive cues to both energize instrumental actions and to selectively guide choice towards actions associated with specific outcomes. Importantly, IC inhibition did not diminish the rats’ motivation to perform an instrumental action in the progressive ratio task and there appeared to be no deficit in the expression of stimulus-outcome associations. Indeed, IC-inactivated rats visited the food port more during the rewarded stimulus than during the unrewarded stimulus in the general transfer experiments, and also visited the food port more frequently during the reward-predictive stimuli than during the baseline period in the specific transfer experiment. However, it is difficult to know whether this food port behaviour reflects an outcome-specific expectancy, as only a single food port was used for delivering the outcomes predicted by both stimuli. Therefore, the Pavlovian conditioned responses observed could result from either a general or specific reward expectancy.

The gustatory region of IC is thus necessary for Pavlovian stimuli to exert both general excitatory and specific influences on instrumental actions, even though Pavlovian conditioned responses and instrumental performance appear to be preserved. The involvement of IC in *both* general and specific transfer is consistent with evidence that IC neurons respond to Pavlovian cues predicting the availability of multiple tastants (Samuelsen et al., 2012, 2013) and show selective responses to auditory cues predicting distinct outcomes (Gardner and Fontanini, 2014; Vincis and Fontanini, 2016; Vincis et al., 2020). Immediate early gene activation is also increased in IC following presentation of a Pavlovian reward-predictive cue (Dardou et al., 2006, 2007; Kerfoot et al., 2007; Saddoris et al., 2009) and this activation appears to be outcome-specific (Saddoris et al., 2009).

Moreover, this anticipatory activity in IC is functionally relevant. Vincis et al (2020) used a perceptual discrimination task in which head-restrained mice were trained to lick a central spout to receive one of four taste cues during a sampling epoch. The mice then had to wait through a short delay before deciding whether to lick left or right for a water reward. Each taste cue signaled that the left or right lick would be rewarded: sweet sucrose or bitter quinine indicated that mice should perform a left lick for the water reward, while sweet maltose or bitter sucrose octaacetate indicated a right lick.

Electrophysiological recordings demonstrated that, within the first 500 ms, IC neurons effectively discriminated between the four tastes. However, during the subsequent delay epoch immediately prior to choice, population decoding of neural activity showed that firing for tastants associated with the same action started to converge. Importantly, while optogenetic inactivation of IC during sampling of the tastant had no impact on performance, the same manipulation during the delay epoch decreased the number of correct choices (Vincis et al., 2020).

Our findings are also in accordance with studies implicating the gustatory region of IC in the retrieval of outcome value in situations where animals must choose between competing actions (Balleine and Dickinson, 2000; Parkes and Balleine, 2013; Parkes et al., 2015, 2018). In these experiments, rats perform two different actions for distinct rewards and then one of the rewards is devalued before a choice test between the two actions. Typically, rats choose the action associated with the non-devalued outcome more than the action associated with the devalued outcome, indicating that they have encoded the new value of the outcome and are able to retrieve this outcome representation to inform their instrumental responding. However, lesions or inactivation of IC renders rats unable to show a preference for the action earning the non-devalued outcome. Specifically, the evidence indicates that the IC is necessary for retrieving the outcome representation (here, the outcome value) (Balleine and Dickinson, 2000; Parkes and Balleine, 2013; Parkes et al., 2015, 2018).

Taken together, these studies demonstrate that the IC is not required to learn about predictive cues or instrumental responses per se but rather to use outcome representations to guide responding (Saddoris et al., 2009). That is, the IC seems to be involved in using associative representations of (taste) outcomes. The associative cybernetic model of instrumental conditioning is currently the most comprehensive theoretical account for transfer effects (Dickinson and Balleine, 1994; Balleine and Ostlund, 2007; Cartoni et al., 2013). A key feature of this model is that transfer is mediated by a stimulus-induced outcome representation, which serves as a stimulus to trigger the representation, and subsequent performance, of the associated action. The emergence of this S^O^-A chain results notably from the ability to use the backward A-O association (i.e., the O-A association), where the representation of the outcome (S^O^) leads to a mental representation of the corresponding action, resulting in the activation of motor responses (S1^O1^-A1; S2^O2^-A2), thus generating specific PIT. In addition, the model proposes that reward predictive stimuli can prompt a general expectancy for reward, which in turn excites motor responses through a S^reward^-A chain.

Thus, the model proposes that both forms of PIT rely on the use of a S-O-A chain through an O-A association (Balleine & Ostlund, 2007). Based on this model, the lack of transfer effects observed in rats with IC inhibition may result from either a failure to represent the outcome as a stimulus (S^O^), which is required to activate the representation of the associated action (A), or an inability to use the triggered A-O representation in a backward manner. In both cases, there appears to be a failure to use the S^O^-A associative chain within S-A memory, reflecting an inability of stimulus-induced outcome expectations to influence instrumental actions. Our results also showed that IC inhibition impaired the selectivity of outcome-induced reinstatement, suggesting a deficit in the ability to use the specific sensory features of the outcome to recall the associated action. Interestingly, non-selective reinstatement remained intact following IC inhibition. That is, IC inhibited rats are not entirely insensitive to the presence of an appetitive outcome. These results appear consistent with outcome devaluation studies showing that IC lesions impair the ability to recall the current outcome value when that outcome is absent but, if the test is conducted under rewarded conditions, these deficits are alleviated (Balleine and Dickinson, 2000). Moreover, IC inhibited rats are also able to correctly reject a devalued outcome when tested in consumption (Parkes and Balleine, 2013; Parkes et al., 2015, 2018).

While we have shown that the IC contributes to the expression of both forms of PIT, distinct insular pathways are likely at play. The IC is anatomically connected to the basolateral amygdala (BLA) (Sripanidkulchai et al., 1984; Yamamoto et al., 1997; Yamamoto, 2006; Gehrlach et al., 2020), the lateral OFC (Barreiros et al., 2021), and the nucleus accumbens shell (Allen et al., 1991), which have all been exclusively implicated in specific PIT. One might therefore hypothesize that the IC-BLA, IC-shell or IC-lateral OFC pathways could support specific, but not general, PIT. Consistent with this suggestion, previous studies have shown that pharmacological inactivation of the BLA reduces the neural excitatory responses to predictive stimuli in the IC (Samuelsen et al., 2012) and evidence from outcome devaluation studies proposes that the BLA updates and encodes information about the value of the instrumental outcome and then sends this information to IC, where it is retrieved to guide behaviour (Parkes & Balleine, 2013).

By contrast, general PIT may require interaction between the IC and the central amygdala (CeA; Gehrlach et al., 2020) and/or NAc core (Wright and Groenewegen, 1996; Reynolds and Zahm, 2005), as these regions are involved in mediating the general excitatory influences of reward-predictive stimuli on instrumental action (Hall et al., 2001; Holland and Gallagher, 2003; Corbit and Balleine, 2005, 2011; Lex and Hauber, 2008; Lingawi and Balleine, 2012). This is supported by the crucial role of CeA in generating stimulus-induced general motivation for rewards (Warlow and Berridge, 2021) and evidence showing the IC-CeA pathway contributes to taste-related choice behaviour (Schiff et al., 2018). We propose that a double dissociation may therefore exist at the cortico-striatal and cortico-amygdalar level, such that IC-core and IC-CeA may support general (but not specific) transfer, and IC-shell and IC-BLA potentially mediate specific (not general) PIT.

Together, our results provide the first demonstration that the IC mediates the ability of predictive cues to exert both a general and specific influence over instrumental responding, likely via the retrieval of outcome representations to guide behaviour.

Moreover, we show that rats with IC inhibition are unable to use the specific sensory features of the outcome to retrieve its associated action. This suggests a deficit in the use of backward A-O associations. These findings significantly contribute to our understanding of the cortical bases of general and specific Pavlovian-instrumental transfer and add to the growing body of literature investigating the broad role of the IC in decision-making.

## Acknowledgements

This work was supported by the French National Agency for Scientific Research (CoCoChoice ANR-19-CE37-0004-07) and the French government in the framework of the University of Bordeaux’s IdEx “Investments for the Future” program/GPR BRAIN_2030. We thank Y. Salafranque for animal care as well as Gilles Courtand and Angélique Faugère for their technical assistance. We also thank Dr. Etienne Coutureau for comments on an earlier version of this manuscript.

